# Cross-sectional volumes and trajectories of the human brain, gray matter, white matter and cerebrospinal fluid in 9,473 typically aging adults

**DOI:** 10.1101/2020.01.12.903658

**Authors:** Andrei Irimia

## Abstract

Accurate knowledge of adult human brain volume (BV) is critical for studies of aging- and disease-related brain alterations, and for monitoring the trajectories of neural and cognitive functions in conditions like Alzheimer’s disease and traumatic brain injury. This scoping meta-analysis aggregates normative reference values for BV and three related volumetrics—gray matter volume (GMV), white matter volume (WMV) and cerebrospinal fluid volume (CSFV)—from typically-aging adults studied cross-sectionally using magnetic resonance imaging (MRI). Drawing from an aggregate sample of 9,473 adults, this study provides (A) regression coefficients *β* describing the age-dependent trajectories of volumetric measures by sex within the range from 20 to 70 years based on both linear and quadratic models, and (B) average values for BV, GMV, WMV and CSFV at the representative ages of 20 (young age), 45 (middle age) and 70 (old age). The results provided synthesize ∼20 years of brain volumetrics research and allow one to estimate BV at any age between 20 and 70. Importantly, however, such estimates should be used and interpreted with caution because they depend on MRI hardware specifications (e.g. scanner manufacturer, magnetic field strength), data acquisition parameters (e.g. spatial resolution, weighting), and brain segmentation algorithms. Guidelines are proposed to facilitate future meta- and mega-analyses of brain volumetrics.

**Disclosure statement:** The author declares that he has no actual or potential conflicts of interest.

## Introduction

The volume of the adult human brain is a fundamental physical descriptor of neuroanatomy (Gazzaniga 2009). For example, in organismic biology, brain volume (BV) provides one of the most tantalizing correlates of neural and behavioral complexity across animal species, including primates (Butler and Hodos 1996). In anthropology, BV and its closely related measure of intracranial volume (ICV) are key criteria for comparing extinct hominid species to one another and to modern humans (Prothero 2007). Studies of human cerebral volumetry and allometry, whether undertaken by neurobiologists, biomedical scientists or anthropologists, frequently rely on accurate knowledge of BV and ICV to compare healthy and diseased brains, and to monitor the temporal trajectories of BV across healthy and diseased populations (C. D. Smith et al. 2007).

BV trajectories are of great interest in healthy aging, in neurodegenerative diseases—e.g. Alzheimer’s disease (AD) and Parkinson’s disease (PD)—but also in other neurological conditions which may accelerate brain atrophy—e.g. traumatic brain injury (TBI) and stroke (Irimia et al. 2014; Irimia et al. 2015; Irimia and Van Horn 2013). Despite numerous past studies which quantified brain volumetrics in health and disease, identifying reliable reference values for these measures is not always straightforward. Importantly, BV varies considerably with age and sex, such that different researchers may require distinct normative statistics for their studies depending on the age and sex compositions of their samples (Cosgrove et al. 2007). Although this variability has been extensively explored and well established (Gur et al. 2002; Rushton and Ankney 1996), it can remain difficult to identify accurate reference statistics for BV due to several important limitations which are shared by many studies. Firstly, researchers may utilize vastly different techniques to assess brain volumetrics (in vivo neuroimaging, post mortem physical measurement, etc.) and their quantitation approaches may differ substantially, which can lead to substantial variability in measurement error across studies (S. M. Smith et al. 2001). Secondly, many reports are based on samples of inadequate sizes and/or compositions, such that the natural variability of brain volumetrics as a function of age and sex has not been quantified satisfactorily. Thirdly, BV and related measures have been reported in a range of formats (tabulations, graphical representations, written narratives, etc.), expressed using a variety of physical units (cubic centimeters, centiliters, etc.) and manipulated algebraically in various ways (e.g., normalized with respect to ICV, averaged across ages or sexes, etc.), such that different studies’ data may require careful and systematic inspection to ensure proper comparison and interpretation. Thus, brain volumetrics reported by different studies can vary substantially even when their statistics are obtained from samples with similar demographics. For these and other reasons, researchers often find it necessary to review the neuroscience literature very thoroughly to identify reports whose methodologies are congruent with one another’s and whose aggregation can provide trustworthy volumetric estimates above and beyond what single studies can afford.

This investigation is a meta-analysis of research studies which utilized in vivo magnetic resonance imaging (MRI) to infer human BV, ICV, gray matter volume (GMV), white matter volume (WMV) and cerebrospinal fluid volume (CSFV) throughout adulthood. Two sets of results are presented: (A) the linear and quadratic regression coefficient for each of these measures as a function of age, across the interval from age 20 to 70, and (B) the values of BV, GMV, WMV and CSFV at the ages of 20 (young age), 45 (middle age) and 70 (old age). Metrics are reported in tabular format to facilitate visual comparison of studies and to provide researchers with the ability to select reports which they deem to be most appropriate for their purposes. For each study listed, numerical values are reported by sex where available, and weighted grand averages are computed to summarize findings across studies. Guidelines are provided to facilitate appropriate use and interpretation of the results by the reader. Finally, recommendations are made to neuroimaging researchers so that future studies of brain volumetrics report results in ways which facilitate meta-analysis and unbiased comparison with other studies. Although neither exhaustive nor definitive, this meta-analysis provides valuable information for a wide range of researchers working in neuroscience, medicine, anthropology, and related fields.

## Methods

### Data sources

Research articles were retrieved via direct search on the ISI Web of Knowledge (webofknowledge.com) and PubMed (https://www.ncbi.nlm.nih.gov/pubmed). The following query syntax was used: (*“brain volume”* OR *“cerebral volume”* OR *“gray matter volume”* OR *“white matter volume”* OR *“intracranial volume”*) AND (*old OR aging OR elderly OR atrophy*). Studies published before 1990 were excluded due to potential drawbacks related to the quality of imaging data and to the paucity of robust volumetric analysis methods prior to that year.

### Study selection

English-language titles and publications were screened manually for relevance. Studies were excluded if they reported incomplete volumetrics data, e.g. if values had not been reported for the entire structures of interest. Data confounded by averaging across both development and adulthood were excluded. Because this study aimed to aggregate volumetrics which are representative of the typical adult population, studies which sampled healthy participants non-randomly were excluded. Such non-random sampling frequently involved inclusion/exclusion criteria which limited the relevance of computed volumetrics to subsets of—rather than to the typical—adult population. For example, studies were excluded if they had specific selection criteria pertaining to factors which could restrict the generalizability of their results, whether related to environmental factors (smoking, alcohol abuse, diet, educational attainment, physical exercise level, etc.) or genetics (e.g. ApoE genetic profile). By contrast, studies which excluded individuals with neurological, psychiatric, and/or metabolic disease were not discarded because such exclusions are necessary to identify healthy adults. Studies were excluded if brain volumetrics had not been normalized by ICV or if normalized values could not be obtained based on the results made available in each study.

### Data extraction and review

Variables were recorded and tabulated separately for men and women, where possible. Literature reports were surveyed to identify two groups of variables of interest:

A. linear regression coefficients {*β*_0_, *β*_1_} and quadratic regression coefficients {*β*_0_, *β*_1_, *β*_2_}, reported as percentages of volumetric differences per year (%/year), for the association between the independent variable (age) and each dependent variable (BV, GMV, WMV or CSFV, respectively);
B. ICV-corrected, cross-sectional average values of BV, GMV, WMV and CSFV at the ages of 20 (young age), 45 (middle age) and 70 (old age), in cm^3^.

A scoping review was conducted using the formalism of Arksey & O’Malley (2005). Scoping reviews are similar to systematic reviews and involve a systematic approach to reference search, thus being less vulnerable to bias compared to rapid, critical or expert reviews (Grant and Booth 2009). By contrast, however, although scoping reviews do not involve systematic quality assessment, they do incorporate more flexible criteria for screening and inclusion.

### Meta-analysis

A meta-analysis was conducted based on an established approach (DerSimonian and Laird 1986, 2015) for aggregating measures of interest over all *i* = 1, …, *M* studies, where *M* is the number of studies included. The meta-analysis was conducted in accordance with PRISMA guidelines (http://www.prisma-statement.org). In the formalism of DerSimonian & Laird, each study’s measured effect *y*_*i*_ is parceled as the sum of the true effect *θ* and the sampling error *e*_*i*_, i.e.

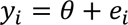

where the variance *σ*^2^ (*e_i_*) is the sample variance 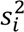 of the effect. To account for the variation in true effects, the model assumes that the true effect is the sum of both *µ* (the mean effect for a population of possible evaluations of the effect), and δ_*i*_ (the deviation of the effect in study *i* from the population’s mean effect *µ*), i.e.

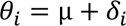

such that

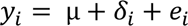

Conceptually, the studies in the meta-analysis are a sample from the population of possible studies evaluating the true effect *θ*, whose population mean is *µ* and whose population variance is *Δ*^2^, such that *θ* ∼ *N*(*µ*, *Δ*^2^) and *yi* ∼ *N*(*θi*, *s_i_*^2^). In the present study, the effect *y_i_* is assumed to be the change in volume *V* observed over some time interval measured in years. Although the regression coefficient is not the most typical measure of effect size, its use as such is not uncommon (Nieminen et al. 2013); our interest in volumetric differences as a function of age makes it very suitable—if not even ideal—for this study.

In meta-analyses where regression coefficients are used to calculate effect size, such coefficients are often standardized to remove the potential confound of them being reported by different studies using different physical units (Nieminen et al. 2013). Here, converting regression coefficients to the same set of physical units (e.g., cm^3^/year) is straightforward; additionally, unstandardized regression coefficients are of substantial interest in practice due to their immediate physical interpretation as BV differences observed over an age interval. For this reason, this analysis utilizes *unstandardized* regression coefficients as measures of effect, after their appropriate conversion to cm^3^/year, where necessary. Because head size may confound brain volumetrics to a substantial extent (Barnes et al. 2010), the effect of this variable is alleviated here by using ICV-normalized volumes—instead of absolute volumes—for all calculations. Thus, the regression coefficients reported here are based on regressions of ICV-normalized volumes rather than on absolute volumes. Furthermore, because scanner field strengths, scanner models, head coil channel number, MRI weightings, sequence parameters and segmentation techniques differed across studies, the potentially confounding effects of these parameters were accounted for by treating the latter as covariates in the meta-analysis provided that their values were available in the original publications. This was accomplished using a weighted least squares technique in which weights were defined by the reciprocal of the variance in the corresponding effect estimate (Sutton et al. 2004; Thompson and Sharp 1999). Study sites were treated as random effects.

Let *β*_1_*_i_* denote the unstandardized linear regression coefficient *β*_1_ associated with study *i*, let 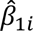 be its empirical estimate, let *β*_0*i*_ and 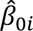 be the intercept and its empirical estimate, respectively, and let *M*_*i*_ be the sample size of study *i*. Of interest here is the meta-analyzed regression coefficient 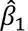, i.e. the slope of the line describing the relationship between time *t* and volume *V*. In the case of a quadratic model, the meta-analyzed regression coefficient 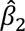 is of additional interest. Without loss of generality, the statistical effect of interest can be assumed to be the volume difference d*V*_*i*_ whose value the regression model of study *i* predicts within some chosen time interval (age range) d*t*. The measured effect *y*_*i*_ defined previously is thus identically equal to d*V*_*i*_ here, i.e. *y*_*i*_ ≡ d*V*_*i*_. In the case of the linear model, 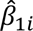 can be computed as d*V*_*i*_/d*t*, where the denominator of the derivative is the change d*t* in chronological age *t*, expressed in units of years and computed identically across all studies (i.e., across the interval from 20 to 70 years of age).

### Heterogeneity analysis

To evaluate effect homogeneity across studies, Cochran’s *Q* statistic

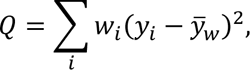

was computed, where

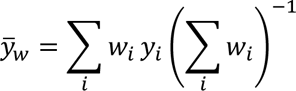

is the weighted estimator of the effect, and *w*_*i*_ is equal to 1/*s*_i_^2^(*y*). Under the null hypothesis, *Q* ∼ *Χ*^2^(*M*_*i*_ − 1). The null hypothesis of homogeneity across studies in the meta-analysis is rejected if *Q* falls within the critical region 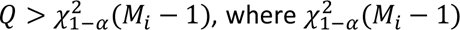, where 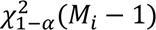 is the (1 − *α*)-quantile of the *Χ*^2^ distribution with *M*_*i*_ − 1 degrees of freedom, and *α* is the significance threshold; in this study, *α* = 0.05. In addition to Cochran’s *Q*, the *I*^2^ index was calculated. This index is set to

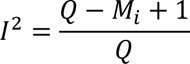

if the right-hand side above is greater than zero and to zero otherwise. *I*^2^ does not depend on the number of studies included in the meta-analysis and can be used to assess consistency between studies. Here, *I*^2^ was interpreted to represent low (*I*^2^ < 0.25), moderate (0.25 ≤ *I*^2^ < 0.50) or high (*I*^2^ ≥ 0.50) inconsistency (Higgins et al. 2003). To identify publication bias, the rank correlations of Begg & Mazumdar (1994) were calculated. Forest plots were generated to visualize results and to identify potential outlier studies.

### Model selection

To investigate the relative merit of linear vs. quadratic models, the Akaike information criterion (AIC) and Bayesian information criterion (BIC) were used. The AIC estimates out-of-sample prediction error and gauges the relative quality of statistical models (Shiavi 2006). The AIC is defined as

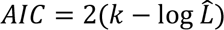

where *k* is the model order and 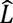 is the empirically-derived maximum value of the model’s likelihood function. In our context, the sampling error *e*_*i*_ can be assumed to be normally distributed and the log-likelihood function assumes the formula

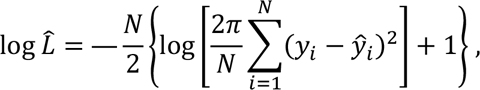

as derived elsewhere (Rossi 2018). When comparing two candidate models *i* and *j*, the quantity

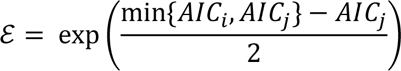

is proportional to the probability that model *j* minimizes the estimated information loss (Burnham and Anderson 2002).

## Results

*Study selection, heterogeneity*, and publication bias. A total of 6,458 published studies were retrieved using the search criteria, of which 5,832 studies were excluded because they were not relevant to the topic of the study. Of the remaining 626 studies, 137 contained incomplete volumetrics data, 57 reported volumetrics which had been averaged over not only adults but also children and/or adolescents, 147 did not study healthy adults who were representative of the typical adult population, 155 reported data on clinical samples rather than healthy adults and 105 reported volumes which had not been normalized by ICV. Notably, many studies which utilized data from the Alzheimer’s Disease Neuroimaging Initiative (ADNI, http://adni.loni.usc.edu), the Parkinson Progression Marker Initiative (PPMI, https://www.ppmi.info.org) or the UK Biobank (https://www.ukbiobank.ac.uk) could not be included for a variety of reasons, such as that (A) their subject sampling methods did not satisfy the inclusion/exclusion criteria of this study (e.g. their sampling of healthy control participants was not fully random), (B) they did not report the numerical quantities of interest here, or (C) the metrics of interest were not reported in a format which could accommodate their accurate analysis using the approach of the present study. In most cases, exclusions due to non-random sampling involved studies whose subjects were substantially healthier or less healthy than in the general population, which is within the scope of this study. Such excluded studies involved individuals who had been selected based on their highly atypical health profile(s) pertaining to cardiovascular, neurovascular, metabolic, neurological, immune, gastrointestinal, urinary, or psychiatric disease. Excluding such subjects was deemed to be useful to avoid producing a meta-analysis which was unlikely to adequately reflect the typical brain volumetrics profiles of the general population. Because such a meta-analysis could have been confounded severely by the highly variable selection criteria of various imaging studies, only studies which sampled randomly from the general population were included despite the likely possibility that some valuable data were thus omitted. After all exclusions, the combined samples of the studies included in the meta-analysis amounted to a total of 9,473 subjects whose volumetrics and related data were aggregated.

Thirty studies were included in the meta-analysis; twelve had fewer than 100 participants (Jernigan et al. 1991; Gur et al. 1991; Matsumae et al. 1996; Guttmann et al. 1998; Jernigan et al. 2001; Ge et al. 2002; Liu et al. 2003; Scahill et al. 2003; Allen et al. 2005; Benedetti et al. 2006; Abe et al. 2008; Barnes et al. 2010), seventeen had between 100 and 1000 participants (Resnick et al. 2000; Good et al. 2001; Gur et al. 2002; Sowell et al. 2003; Taki et al. 2004; Fotenos et al. 2005; Lemaitre et al. 2005; Chen et al. 2007; C. D. Smith et al. 2007; Fotenos et al. 2008; Driscoll et al. 2009; Michielse et al. 2010; Walhovd et al. 2011; Lemaitre et al. 2012; Peelle et al. 2012; Jancke et al. 2015; Blatter et al. 1995) and one had over 1000 participants (DeCarli et al. 2005). Studies were not found to be significantly heterogeneous when reporting BV (*Q* = 0.29, *p* > 0.99, *I*^2^ = 0), GMV (*Q* = 1.15, *p* > 0.99, *I*^2^ = 0) or WMV (*Q* = 1.35, *p* > 0.99, *I*^2^ = 0), but were found to have mild heterogeneity when reporting CSFV (*Q* = 15.49, *p* > 0.22, *I*^2^ = 0.23). The Begg-Mazumdar rank correlation (*τ* = 0.12, *p* > 0.67) did not provide significant indication of publication bias. The results of the meta-analysis are summarized in Table 1, Table 2 and Table 3, which contain (A) 9 studies reported only in Table 1; (B) 7 studies reported in both Table 1 and Table 2; (C) 9 studies reported only in Table 3, (D) 5 studies reported in both Table 1 and Table 3; (E) 12 studies reported in both Table 1 and Table 3, and (F) 5 studies reported in Table 1, Table 2 and Table 3, for a total of 30 studies in the entire meta-analysis. Table 4 reports study parameters for all 30 studies. Figure 1 reports information based on all studies listed in Table 1.

**Figure 1.**
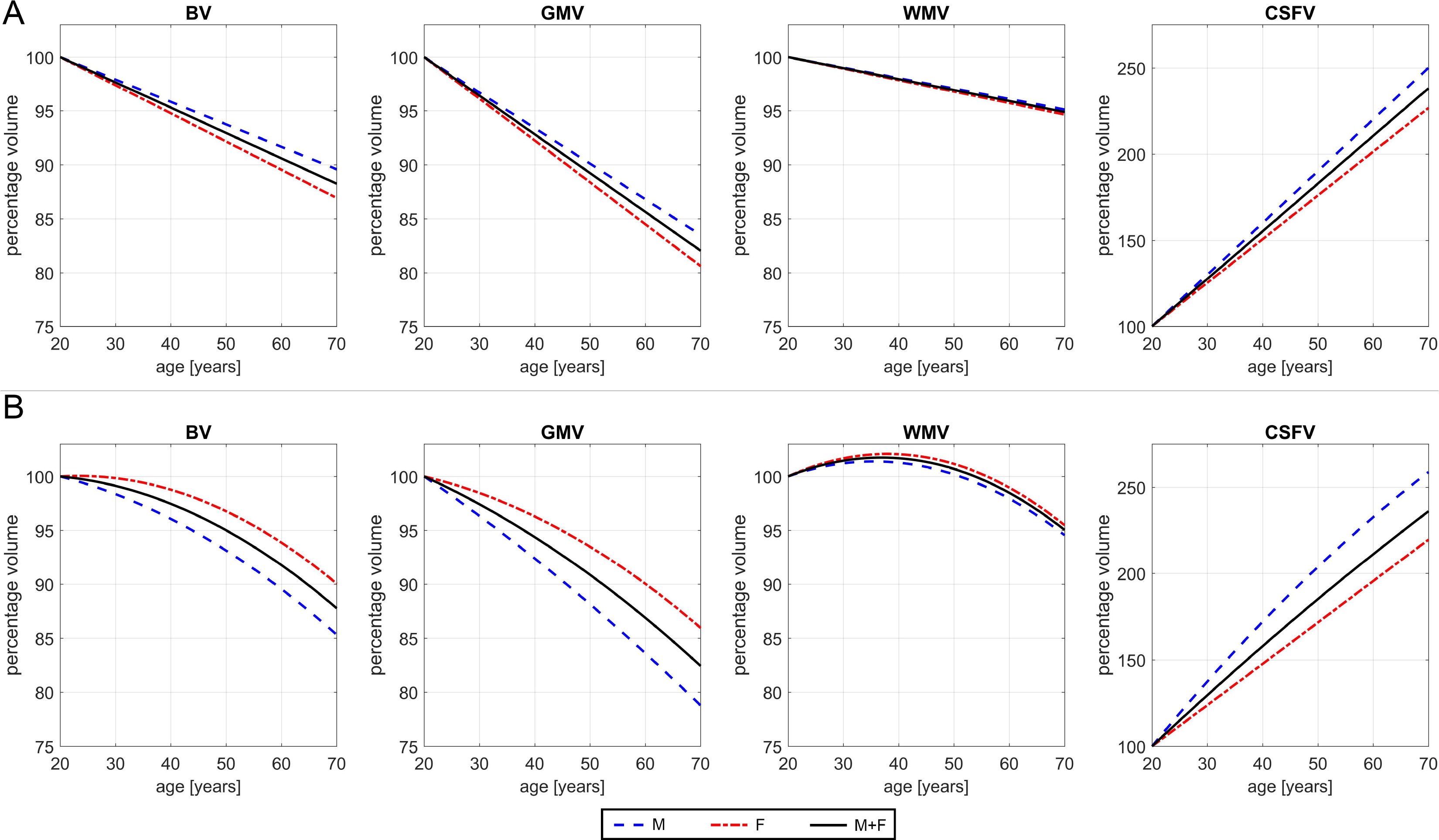
Forest plots of regression coefficients (A) and volumes (B) for the brain (B, first column), gray matter (GM, second column), white matter (WM, third column) and cerebrospinal fluid (CSF, fourth column). The first author of each study included in the meta-analysis is listed on the left; studies are listed in ascending chronological order, with the oldest one at the top and the newest one at the bottom. Means are indicates by red squares whose sizes are proportional to studies’ sample sizes; standard deviations are indicated by horizontal lines. In (A), the regression coefficients *β* are expressed as percentages of volume difference per year [%/year]. In (B), volumes are expressed in cm^3^. See Tables 1 for additional data.

**Table 1.**
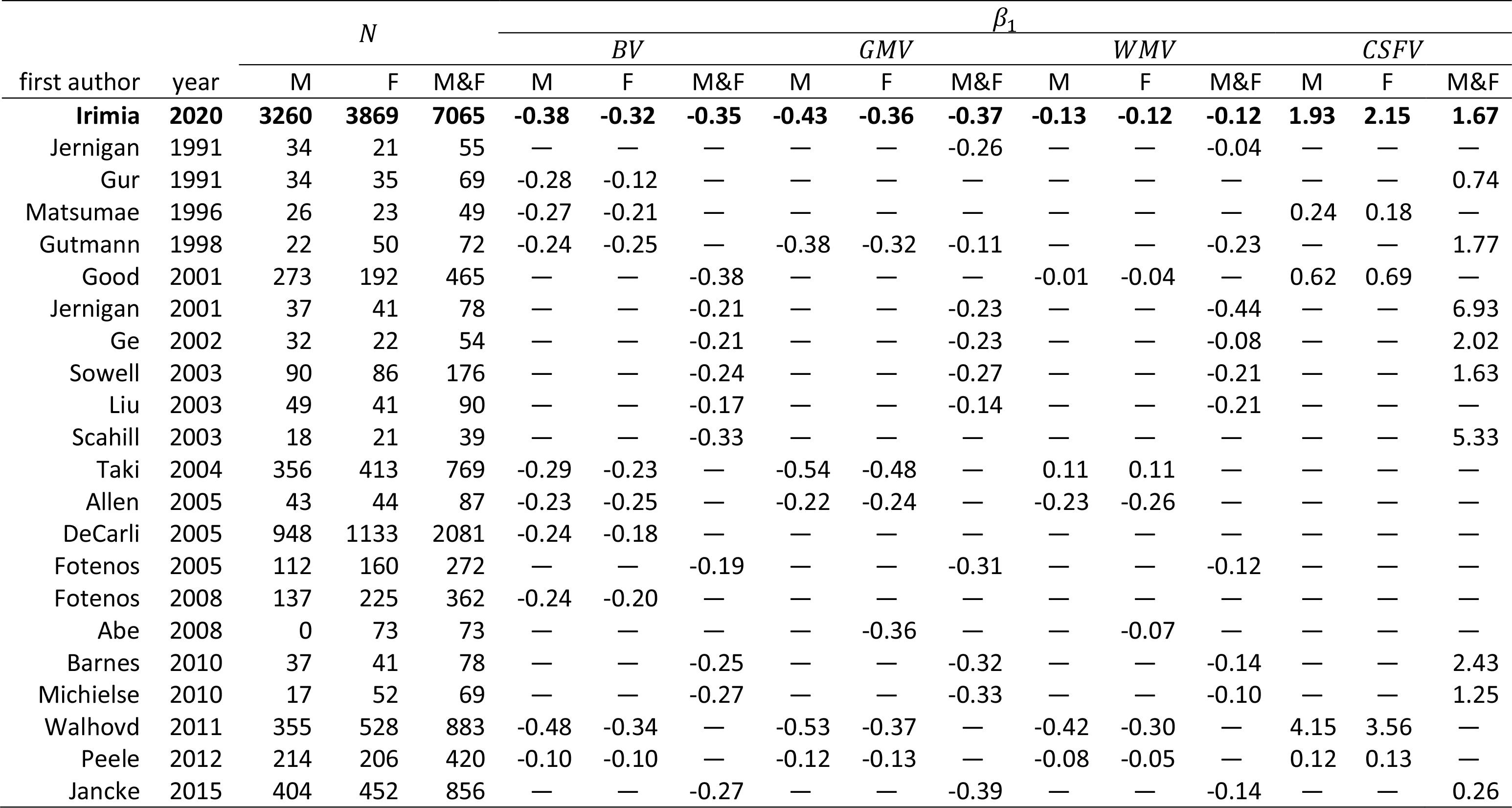
Linear regression coefficients *β*_*i*_ conveying the annualized change in the ICV-normalized BV, GMV, WMV, and CSFV. Regression coefficients are reported as percentage differences in volume per year (%/year). Meta-analysis results are displayed in bold face. Regression coefficients are listed separately for males and females, where possible; for studies which do not report regression coefficients separately for males (M) and females (F), regression coefficients are listed under M&F. All linear regression models apply to the age interval between 20 and 70 years and assume that volumetric changes occur linearly within this interval. The sample size *N* of each study is also reported. A dash indicates that data were not reported by the study in question, or that reported data could not be utilized in the meta-analysis for methodological reasons (see text). Abbreviations: M = males; F = females; M&F = males and females; BV = brain volume; GMV = gray matter volume; WMV = white matter volume; CSFV = cerebrospinal fluid volume.

**Table 2.** Quadratic regression coefficients (*β*_0_, *β*_1_, *β*_2_), their confidence intervals (CIs) and their pooled, empirical standard deviations (*s*) for BV, GMV, WMV and CSFV. The aggregate sample contains 3,897 subjects pooled across 7 studies where quadratic fits were reported (see text for details). Like in Table 1, the regression coefficients are reported as percentage differences in volume per year (%/year) and convey the annualized change in the ICV-normalized BV, GMV, WMV and CSFV. Abbreviations: BV = brain volume; GMV = gray matter volume; WMV = white matter volume; CSFV = cerebrospinal fluid volume; CI = confidence interval.

| metric | $\beta_0$ | $s(\beta_0)$ | CI ($\beta_0$) | $\beta_1$ | $s(\beta_1)$ | CI ($\beta_1$) | $\beta_2$ | CI ($\beta_2$) | $s(\beta_2)$ |
| --- | --- | --- | --- | --- | --- | --- | --- | --- | --- |
| BV | 99.34 | 0.89 | [ 97.56, 101.12] | 0.11 | 0.01 | [ 0.09, 0.13] | -0.004 | [-0.005, -0.003] | 0.0012 |
| GMV | 103.93 | 2.57 | [101.36, 106.50] | -0.14 | 0.01 | [-0.16, -0.12] | -0.002 | [-0.003, -0.001] | 0.0011 |
| WMV | 93.43 | 1.29 | [ 90.19, 96.67] | 0.45 | 0.01 | [ 0.43, 0.47] | -0.006 | [-0.008, -0.004] | 0.0011 |
| CSFV | 37.61 | 2.46 | [ 32.69, 42.53] | 3.23 | 0.18 | [ 2.87, 3.59] | -0.006 | [-0.007, -0.005] | 0.0005 |

**Table 3.** Average ICV, BV, GMV, WMV and CSFV at the ages of 20, 45 and 70. Meta-analysis results which aggregate all studies listed in the table are displayed in ace. Where possible, values are listed separately for males and females; for studies which do not distinguish trajectories by sex, values are listed under M&F. mple size *N* of each study is also reported. A dash indicates that data were not reported by the study in question, or that reported data could not be utilized meta-analysis for methodological reasons (see text). Abbreviations: M = males; F = females; M&F = males and females; ICV = intracranial volume; BV = brain e; GMV = gray matter volume; WMV = white matter volume; CSFV = cerebrospinal fluid volume.

|  |  | first author | Irimia | Gur | Blatter | Resnick | Good | Gur | Sowell | Scahill | Taki | Allen | DeCarli | Lemaître | Benedetti | Chen | Smith | Abe | Fotenos | Driscoll | Michielse | Walhovd | Lemaître | Peele |
| --- | --- | --- | --- | --- | --- | --- | --- | --- | --- | --- | --- | --- | --- | --- | --- | --- | --- | --- | --- | --- | --- | --- | --- | --- |
|  | age | year | 2020 | 1991 | 1995 | 2000 | 2001 | 2002 | 2003 | 2003 | 2004 | 2005 | 2005 | 2005 | 2006 | 2007 | 2007 | 2008 | 2008 | 2009 | 2010 | 2011 | 2012 | 2012 |
| N | all | M | **3474** | 34 | 89 | 68 | 273 | 57 | 90 | 18 | 356 | 43 | 948 | 331 | 39 | 184 | 51 | 0 | 137 | 73 | 17 | 355 | 97 | 214 |
|  |  | F | **4065** | 35 | 105 | 48 | 192 | 59 | 86 | 21 | 413 | 44 | 1133 | 331 | 50 | 227 | 71 | 73 | 225 | 47 | 52 | 528 | 119 | 206 |
|  |  | M&F | **7539** | 69 | 194 | 116 | 465 | 116 | 176 | 39 | 769 | 87 | 2081 | 662 | 89 | 411 | 122 | 73 | 362 | 120 | 69 | 883 | 216 | 420 |
| ICV | 20 | M | **1533** | — | 1594 | — | 1690 | — | — | — | — | — | 1400 | — | — | — | — | — | — | — | — | 1694 | — | 1625 |
|  |  | F | **1374** | — | 1400 | — | 1550 | — | — | — | — | — | 1250 | — | — | — | — | 1450 | — | — | — | 1510 | — | 1500 |
|  |  | M&F | **1446** | — | — | — | — | — | 1450 | — | — | — | — | — | — | — | — | — | — | — | 1400 | — | — | — |
|  | 45 | M | **1516** | — | 1538 | — | 1640 | — | — | — | — | — | 1360 | — | — | 1650 | — | — | — | — | — | 1695 | — | 1625 |
|  |  | F | **1358** | — | 1347 | — | 1540 | — | — | — | — | — | 1210 | — | — | 1440 | — | 1438 | — | — | — | 1531 | — | 1450 |
|  |  | M&F | **1430** | — | — | — | — | — | 1445 | 1390 | — | — | — | — | — | — | — | — | — | — | 1400 | — | — | — |
|  | 70 | M | **1434** | — | 1548 | — | 1630 | — | — | — | — | 997 | 1320 | — | — | — | — | — | — | — | — | 1662 | — | 1620 |
|  |  | F | **1296** | — | 1335 | — | 1530 | — | — | — | — | 882 | 1150 | — | — | — | — | 1425 | — | — | — | 1469 | — | 1450 |
|  |  | M&F | **1373** | — | — | — | — | — | 1390 | 1390 | — | — | — | — | — | — | — | — | — | — | 1400 | — | — | — |
| BV | 20 | M | **1185** | 1180 | 1368 | — | 1315 | 1240 | — | — | — | 1123 | 1100 | — | — | — | — | — | — | — | — | 1176 | — | 1330 |
|  |  | F | **1110** | 1090 | 1354 | — | 1175 | 1210 | — | — | — | 1009 | 1050 | — | — | — | — | 1143 | — | — | — | 1119 | — | 1220 |
|  |  | M&F | **1149** | — | — | — | — | — | 1310 | — | — | — | — | — | — | — | — | — | 1190 | — | 832 | — | — | — |
|  | 45 | M | **1179** | 1140 | 1316 | — | 1225 | 1190 | — | — | — | 1059 | 1100 | — | — | — | — | — | — | — | — | 1247 | — | 1330 |
|  |  | F | **1051** | 1050 | 1323 | — | 1125 | 1070 | — | — | — | 943 | 980 | — | — | — | — | 1043 | — | — | — | 1088 | — | 1160 |
|  |  | M&F | **1116** | — | — | — | — | — | 1255 | 1240 | — | — | — | — | — | — | — | — | 1160 | — | 800 | — | — | — |
|  | 70 | M | **1067** | 1080 | 1268 | 1130 | 1125 | — | — | — | — | 920 | 1003 | 1066 | — | — | — | — | — | — | — | 1030 | — | 1260 |
|  |  | F | **947** | 1050 | 1285 | 920 | 1025 | — | — | — | — | 790 | 874 | 960 | — | — | — | 988 | — | — | — | 929 | — | 1140 |
|  |  | M&F | **1009** | — | — | — | — | — | 1150 | 1140 | — | — | — | — | — | — | — | — | 1080 | 950 | 725 | — | — | — |
| GMV | 20 | M | **749** | — | 689 | — | 865 | 710 | — | — | 700 | 604 | — | — | — | — | — | — | — | — | — | 728 | — | 780 |
|  |  | F | **658** | — | 638 | — | 785 | 660 | — | — | 590 | 555 | — | — | — | — | — | 730 | — | — | — | 644 | — | 720 |
|  |  | M&F | **692** | — | — | — | — | — | 830 | — | — | — | — | — | 790 | — | — | — | — | — | 520 | — | 495 | — |
|  | 45 | M | **694** | — | 682 | — | 775 | 660 | — | — | 600 | 571 | — | — | — | 741 | — | — | — | — | — | 684 | — | 760 |
|  |  | F | **612** | — | 640 | — | 715 | 610 | — | — | 520 | 521 | — | — | — | 667 | — | 630 | — | — | — | 600 | — | 670 |
|  |  | M&F | **641** | — | — | — | — | — | 730 | — | — | — | — | — | 760 | — | — | — | — | — | 475 | — | 440 | — |
|  | 70 | M | **598** | — | 644 | 570 | 685 | — | — | — | 520 | 460 | — | 575 | — | — | — | — | — | — | — | 562 | — | 730 |
|  |  | F | **540** | — | 648 | 500 | 650 | — | — | — | 470 | 395 | — | 532 | — | — | — | 575 | — | — | — | 507 | — | 650 |
|  |  | M&F | **560** | — | — | — | — | — | 680 | — | — | — | — | — | — | — | 560 | — | — | 525 | 425 | — | 400 | — |
| WMV | 20 | M | **509** | — | 678 | — | 450 | 530 | — | — | 460 | 519 | — | — | — | — | — | — | — | — | — | 531 | — | 550 |
|  |  | F | **460** | — | 638 | — | 390 | 450 | — | — | 420 | 454 | — | — | — | — | — | 413 | — | — | — | 475 | — | 500 |
|  |  | M&F | **488** | — | — | — | — | — | 480 | — | — | — | — | — | 770 | — | — | — | — | — | 312 | — | — | — |
|  | 45 | M | **518** | — | 633 | — | 450 | 570 | — | — | 470 | 488 | — | — | — | 496 | — | — | — | — | — | 563 | — | 570 |
|  |  | F | **464** | — | 683 | — | 400 | 460 | — | — | 430 | 422 | — | — | — | 425 | — | 413 | — | — | — | 488 | — | 490 |
|  |  | M&F | **494** | — | — | — | — | — | 525 | — | — | — | — | — | 710 | — | — | — | — | — | 325 | — | — | — |
|  | 70 | M | **486** | — | 624 | 460 | 440 | — | — | — | 480 | 460 | — | 491 | — | — | — | — | — | — | — | 468 | — | 530 |
|  |  | F | **442** | — | 638 | 420 | 375 | — | — | — | 450 | 395 | — | 428 | — | — | — | 413 | — | — | — | 421 | — | 490 |
|  |  | M&F | **457** | — | — | — | — | — | 470 | — | — | — | — | — | — | — | 380 | — | — | 425 | 300 | — | — | — |
| CSFV | 20 | M | **188** | 80 | 78 | — | 370 | 95 | — | — | — | — | — | — | — | — | — | — | — | — | — | 20 | — | 320 |
|  |  | F | **136** | 75 | 91 | — | 370 | 80 | — | — | — | — | — | — | — | — | — | — | — | — | — | 16 | — | 275 |
|  |  | M&F | **159** | — | — | — | — | — | 140 | — | — | — | — | — | — | — | — | — | — | — | 160 | — | — | — |
|  | 45 | M | **300** | 140 | 129 | — | 420 | 110 | — | — | — | — | — | — | — | 410 | — | — | — | — | — | 224 | — | 325 |
|  |  | F | **253** | 100 | 122 | — | 410 | 100 | — | — | — | — | — | — | — | 350 | — | — | — | — | — | 196 | — | 280 |
|  |  | M&F | **266** | — | — | — | — | — | 190 | 150 | — | — | — | — | — | — | — | — | — | — | 180 | — | — | — |
|  | 70 | M | **413** | 210 | 178 | — | 480 | — | — | — | — | — | — | — | — | — | — | — | — | — | — | 489 | — | 330 |
|  |  | F | **352** | 160 | 165 | — | 500 | — | — | — | — | — | — | — | — | — | — | — | — | — | — | 372 | — | 290 |
|  |  | M&F | **365** | — | — | — | — | — | 240 | 250 | — | — | — | — | — | — | 390 | — | — | — | 255 | — | — | — |

**Table 4.**
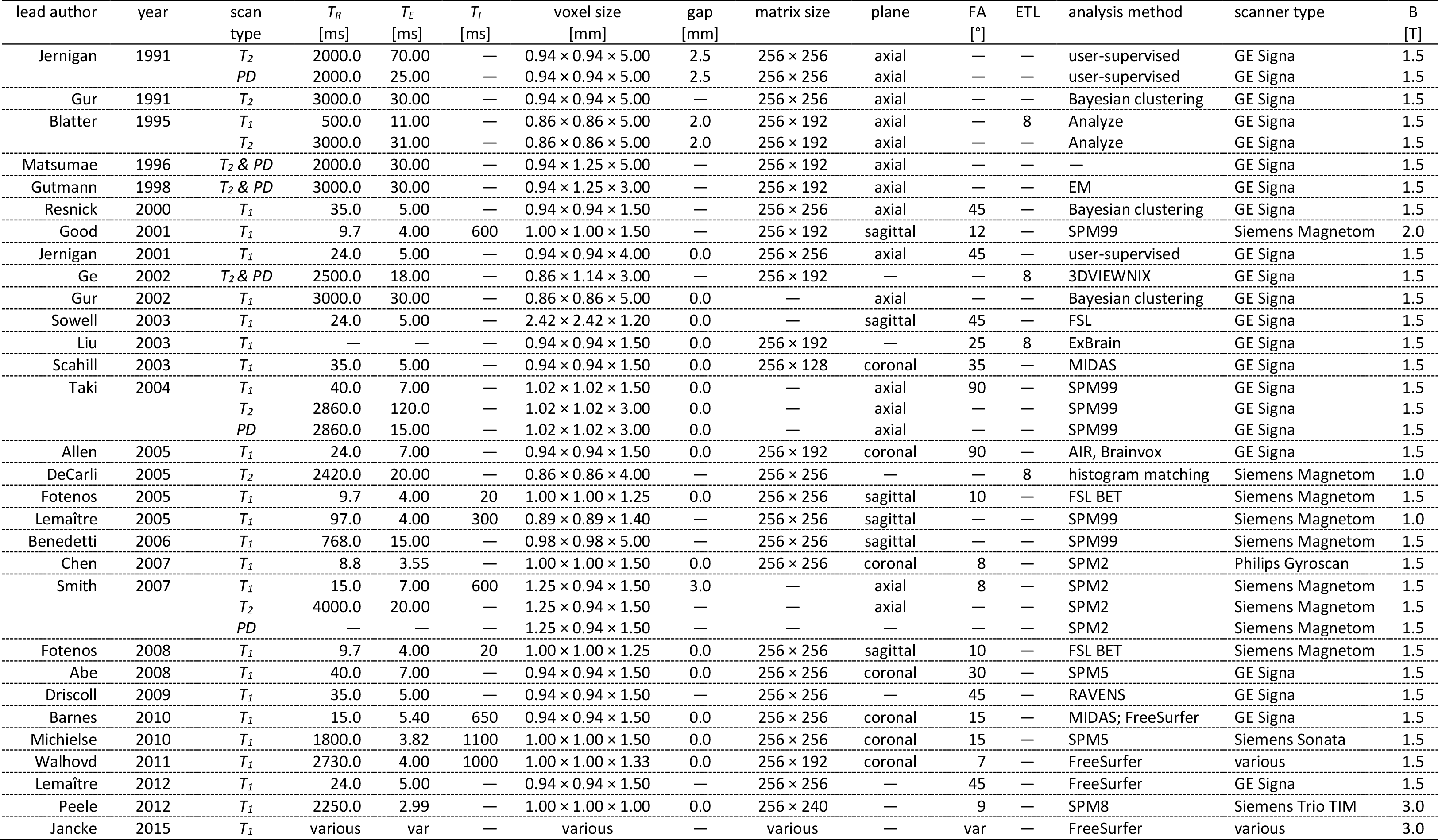
MRI acquisition parameters for the studies included in the meta-analysis. Dashes indicate either that the parameter in question was not specified in the original publication or that it does not apply to the protocol used in the study. If a software package was used for analysis, this is indicated. If an in-house method was used, the type of segmentation method is stated. Notes: In the last row, the study of Jancke et al. includes data from 16 distinct *T_1_*-weighted samples acquired at 3 T using various parameters; this is indicated as *var* or *various*; however, all data were analyzed using FreeSurfer. For the 1996 study of Matsumae et al. and the 1998 study of Gutman et al., *T_2_*-weighted and PD images were acquired on the same scanner using interleaved sequences with identical parameters. For the 2002 study by Ge et al., PD images were utilized but their acquisition parameters are not stated in the original publication. Abbreviations: AIR = automated image registration; B = magnetic field strength; BET = brain extraction tool; EM = expectation maximization; ETL = echo train length; ExBrain = extended brain; FSL = FMRIB software library; GE = General Electric; MIDAS = metabolite imaging and data analysis system; mm = millimeter; ms = millisecond; PD = proton density; RAVENS = regional analysis of volumes examined in normalized space; SPM = statistical parametric mapping; T = tesla; *T_E_* = echo time; *T_I_* = inversion time; *T_R_* = repetition time; var = various.

### Linear regression coefficients

Table 1 and Figure 1 (A) summarize the cross-sectional linear trajectories of BV, GMV, WMV and CSFV as reported by studies fitting the selection criteria of the analysis. The regression coefficients (slopes) for volume differences as a function of age are reported as percentages of volumetric difference per year (%/year) and are listed separately for males (M) and females (F), where available. For studies where numbers are reported for combined samples, the numerical values in question are listed in the column labeled as “M&F.” Studies are listed in ascending order by year of publication, and their sample sizes are reported. In all cases, 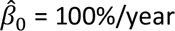 for these linear models and is therefore not reported in Table 1. For BV, the estimated values of the linear regression coefficient 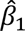, pooled standard deviation *s* and confidence interval (CI) calculated across the entire aggregate sample was found to be 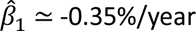 (pooled 2*s* = 0.05%/year; 95% CI = [-0.40, -0.30] %/year). For GMV, 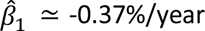 (pooled 2*s* = 0.05%/year; 95% CI = [-0.42, -0.32] %/year); for WMV, 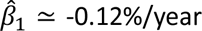 (pooled 2*s* = 0.012%/year; 95% CI = [-0.13, -0.11] %/year); for CSFV, 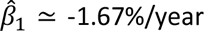 (pooled 2*s* = 0.47%/year; 95% CI = [-2.14, -1.12] %/year). The pooled value of twice the standard deviation *s* at age ∼20 was 0.043%/year for BV, 0.045%/year for GMV, 0.008%/year for WMV and 0.42%/year for CSFV. At age ∼45, it was 0.051%/year for BV, 0.052%/year for GMV, 0.010%/year for WMV and 0.48%/year for CSFV. At age ∼70, it was 0.057%/year for BV, 0.055%/year for GMV, 0.012%/year for WMV and 0.54%/year for CSFV.

Table 1 includes, in boldface, the regression coefficients for each aggregate sample. Specifically, the average year-to-year difference in BV was found to be -0.35% across sexes (males: -0.38%; females: - 0.32%); studies reported values of 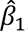 ranging from about -0.40%/year (Walhovd et al. 2011) to about - 0.10%/year (Peelle et al. 2012). GMV was found to trend negatively with age (males: -0.43%; females: - 0.36%; both sexes: -0.37%), and more so than WMV (males: -0.13%; females: -0.12%; both sexes: -0.12%). The annual difference in CSFV was found to be smaller in males (1.93%) than in females (2.15%), although the computed difference is relatively small (0.22%). Furthermore, because both GMV and WMV were found to have more negative regression coefficients in males than in females, it is likely that the true slope of the CSFV trajectory as a function of age is steeper in males than in females. Table 1 suggests that the small difference observed between males and females is likely driven by the study of Good et al. (2001), who reported a smaller regression coefficient for males than for females and whose sample size was relatively large compared to most of the other studies included.

Cross-sectionally, over the interval from age 20 to age 70, all studies reported negative trajectories for BV, GMV and WMV; only Taki et al. (2004) reported a positive regression coefficient for WMV. Furthermore, as expected, all studies reported increases in CSFV. These results are confirmed by and reflected in the forest plots of Figure 1 (A). It is important to note that values reported under “M&F” should not be interpreted as weighted averages of the values reported for each of the two sexes. Rather, they represent volumetrics reported by studies where results were not calculated separately for males and females.

### Quadratic regression coefficients

Seven studies (Ge et al. 2002; Sowell et al. 2003; DeCarli et al. 2005; Fotenos et al. 2005; Fotenos et al. 2008; Michielse et al. 2010; Walhovd et al. 2011) reported quadratic model regression coefficients; this aggregate sample includes *N* = 3,897 subjects. The quadratic regression coefficients {*β*_0_, *β*_1_, *β*_2_ } for BV, GMV, WMV and CSFV are reported in Table 2, together with their CIs and their pooled, empirical standard deviations *s*. Both within and across sexes, Figure 2 allows the reader to compare the linear and quadratic regression models of the meta-analysis for BV, GMV, WMV and CSFV. Quadratic models capture the relatively shallower slopes of BV and GMV trajectories during the third and fourth decade of life, followed by steeper slopes thereafter, in agreement with the findings of the meta-analysis (see Table 3). Quadratic models additionally capture the relatively slight overshoot of WMV during middle age relative to its baseline level; this effect is well documented by many of the studies meta-analyzed here. For CSFV, visual inspection of Figure 2 suggests that the two models differ relatively little. In all cases, females’ volumes have trajectories with shallower slopes compared to those of males, reflecting meta-analysis findings (see Table 3).

**Figure 2.**
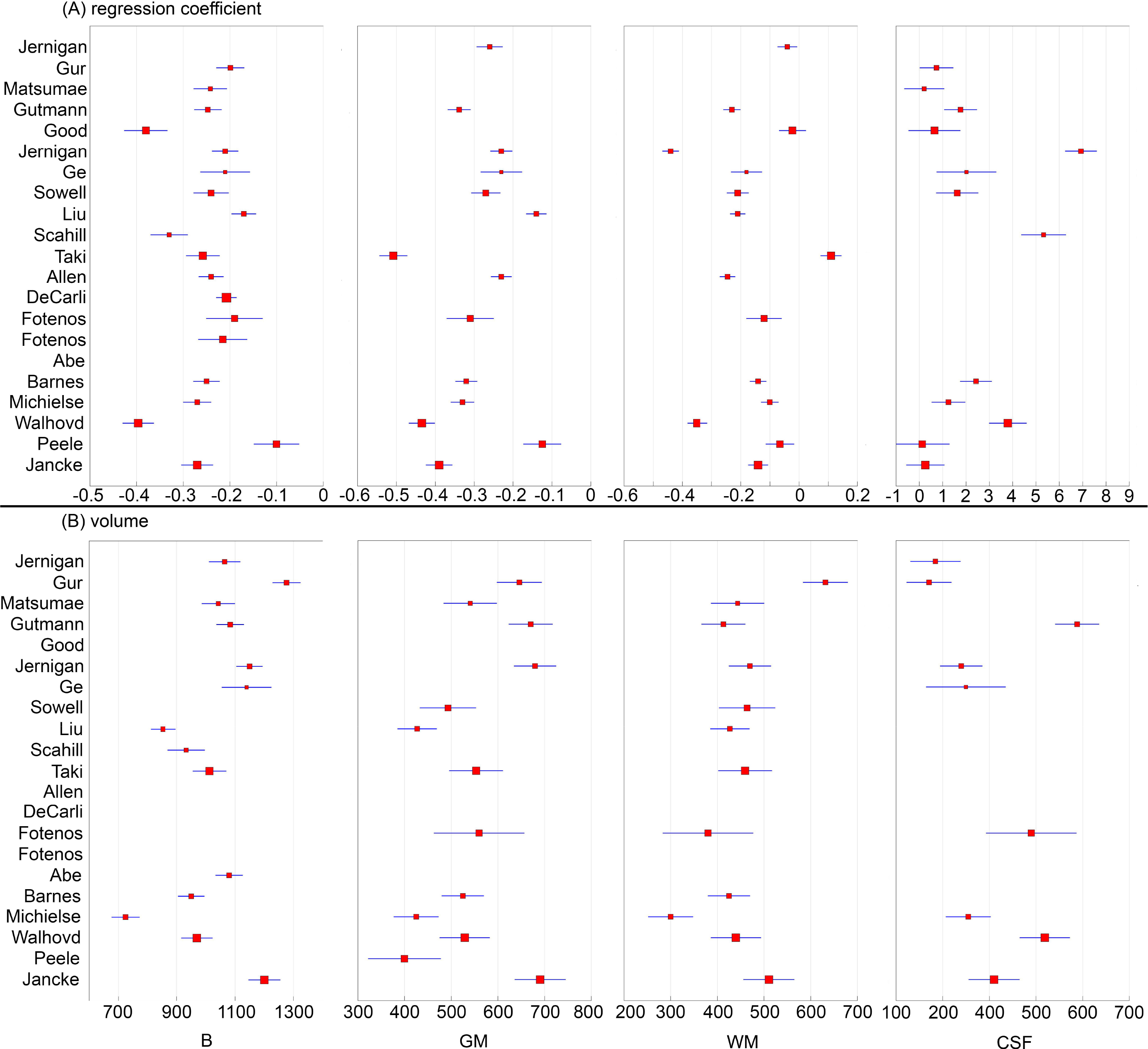
Age-dependent trajectories of brain volume (BV), gray matter volume (GMV), white matter volume (WMV) and CSF volume (CSFV) derived meta-analytically. The top and bottom rows display linear (A) and quadratic (B) models, respectively. The quantity on the vertical axis is the percentage volume, assuming 100% volume at age 20. The quantity on the horizontal axis is age in years. Traces for males and females are shown in red (dashed and dotted lines) and blue (dashed lines), respectively. Traces for both sexes are drawn in black (continuous lines). Because the curves displayed here are cross-sectional volumetric trajectories based on linear and quadratic models, any biological aging effects unaccounted for by such models are not captured by these visual representations.

### Model selection

The quantity ℰ was found to be equal to 0.916 for BV, indicating that the linear model is 91.6% as likely as the quadratic model to minimize information loss for this measure. For GMV, ℰ = 0.842, suggesting that the quadratic model is 84.2% as likely to minimize information loss as the linear model. For WMV and CSFV, the percentages found were 65.7% and 93.3%, respectively.

Average volumetrics. Studies reporting volumetrics were not found to be significantly heterogeneous (*Q* = 16.11, *p* > 0.71, *I*^2^ = 0). The Begg-Mazumdar rank correlation (*τ* = 0.09, *p* > 0.70) failed to provide significant indication of publication bias. Table 3 reports the average volumes of BV, GMV, WMV and CSFV at the ages of 20, 45 and 70, as calculated over studies fitting the selection criteria of the analysis; for brevity, Figure 1 (B) reports this information only for age 70. Volumes are reported in cm^3^ for males (M) and females (F), where available. For studies where numbers are reported for combined samples, the figures in question are listed in the column labeled as “M&F.” Studies are listed in ascending order by year of publication, and their sample sizes are also reported. Table 3 also reports the results of the meta-analysis, i.e. average volumes aggregated across all the studies and samples listed in the table.

ICV was found to decrease very slightly as a function of age (aggregate mean at age 20: ∼1,446 cm^3^; aggregate mean at age 45: ∼1,430 cm^3^; aggregate mean at age 70: ∼1,373 cm^3^), and Table 3 indicates that this finding is consistent across studies. This effect was previously reported elsewhere but its causes remain uncertain (Jantz and Jantz 2016; Jantz and Meadows Jantz 2000). Average BV is found to vary from ∼1,150 cm^3^ at age 20 to ∼1,116 cm^3^ at age 45 and ∼1,009 cm^3^ at age 70. Similarly, GMV varies from ∼692 cm^3^ at age 20 to ∼641 cm^3^ at age 45 and to ∼560 cm^3^ at age 70. Average WMV is ∼509 cm^3^ at age 20, ∼494 cm^3^ at age 45 and ∼457 cm^3^ at age 70. As expected, the trend is reversed for CSFV, which varies from ∼159 cm^3^ at age 20 to ∼266 cm^3^ at age 45 and to ∼365 cm^3^ at age 70. Consistently, average BV, GMV and WMV are larger for males than for females; the reverse is found for CSFV, as expected.

### Key studies

Several of the studies included in this meta-analysis deserve discussion due to their relatively large sample sizes or to their detailed study of volumetrics. One such study is that of Good et al. (2001), who used voxel-based morphometry (VBM) and statistical parametric mapping (SPM) to calculate the brain volumetrics of 465 healthy adults aged 17-79 (265 males) based on *T*_1_-weighted MRIs (voxel size: 1 mm × 1 mm × 1.5 mm). These authors found a negative trend of global GMV with age 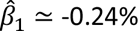 per year across sexes, *R*^2^ = 0.489, *p* < 0.0001), and a GMV/WMV fractional volume ratio of ∼1.82 across sexes.

For CSFV, the ICV-normalized regression coefficient 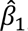 was positive, consistent with age-related increases in ventricular volume (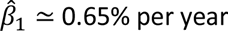, *R*^2^ = 0.377, *p* < 0.0001). Overall, BV was found to decrease in both sexes (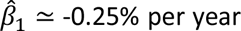, *R*^2^ = 0.489, *p* < 0.0001). Usefully, Good et al. reported both absolute and ICV-corrected values, such that the confound of head size was accounted for in their analysis. Nevertheless, their study had considerably more volunteers younger than 40, such that their results might reflect younger adults’ volumetrics to a greater extent than other studies. Had the authors’ sample been more balanced, the reported regression coefficients would have likely been more negative for GMV, WMV and BV, and more positive for CSFV. Thus, the study of Good et al. may be particularly amenable for comparison to studies involving individuals under the age of 40 who were imaged at today’s standard spatial resolution for MRI.

Taki et al. (2004) used SPM to extract GMVs and WMVs from the *T*_1_-weighted MRIs (voxel size: 1.02 mm × 1.02 mm × 1.5 mm) of 769 Japanese adults aged 16-79 (356 males). Age-related declines were found in global GM (males: 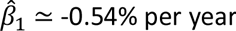, *R*^2^ = 0.58, *p* < 001; females: 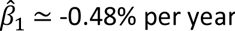, *R*^2^ = 0.39, *p* < 001), but WMV was found to increase with age, albeit only slightly (males: 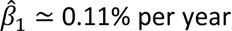, *R*^2^ = 0.017, *p* < 001; females: 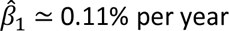, *R*^2^ = 0.019, *p* < 001). For GMV, the age-related volumetric decline rates reported by Taki et al. are relatively faster than in most other studies included in this meta-analysis, whereas for WMV the rates are slightly positive. One limitation of the study by Taki et al. is that it included only adults of Japanese ethnicity, although this can also be perceived as a strength because it allows one to compare their results to those obtained from cohorts of other ethnicities; thus, the study of Taki et al. may be useful in studies of East Asian populations.

DeCarli et al. (2005) acquired *T*_2_-weighted MRIs (field of view: 22 cm; acquisition matrix size: 182 × 256) from 2,081 adults enrolled in the Framingham Heart Study. In this study, regression coefficients were found to be negative, and similar to those of other studies reviewed (males: 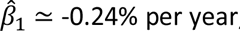, *R*^2^ = 0.45, *p* < 001; females: 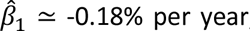, *R*^2^ = 0.35, *p* < 001). GMV and WMV trajectories were reported by lobe, but global statistics were not provided. Although the sample consisted mostly of European Americans—which can be a limitation—this is the largest study included in the meta-analysis. Despite their rigorous study, the results of DeCarli et al. may warrant cautious interpretation and generalization because their MRI data were acquired at relatively low field strength (1.0 T) and spatial resolution (4 mm slice thickness).

Walhovd et al. (2011) combined one Swedish, two Norwegian and three US cohorts to study GMV, WMV, CSFV and BV in an aggregate sample of 883 adults aged 18-94 (355 males). This very thorough study reported not only the regression coefficients of interest here, but also the mean and standard deviations of volumes for CSF structures, WM as well as for cortical and subcortical GM. Volumetric trajectories were reported both by decade and across the entire sample. As in most other studies included in the present analysis, the authors found negative regression coefficients for BV (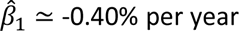, *R*^2^ = 0.47, *p* < 0.001), cerebral GMV (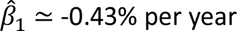, *R*^2^ = 0.54, *p* < 0.001), cerebral WMV (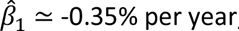, *R*^2^ = 0.12, *p* < 0.001) and ventricular CSFV (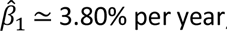, *R*^2^ = 0.37, *p* < 0.001). Compared to many other studies included in this meta-analysis, that of Walhovd et al. stands out because it reports systematic volumetrics in considerable detail. Nevertheless, the results of these authors warrant further confirmation and comparison to other populations partly because the sample included a large proportion of Northern Europeans and because the cross-sectional volumetric trajectories reported are considerably steeper and more negative than in many other studies included in this meta-analysis. Furthermore, there are substantial discrepancies in mean educational attainment between the Northern Europeans and the US participants included, which adds to the sample’s heterogeneity above and beyond other potential confounds related to imaging protocols across sites.

## Discussion

### Motivation and rationale

Identifying factors which influence brain senescence and atrophy and quantifying the extent to which these factors affect brain volume loss are key to understanding the aging process. The relationship between volumetrics and chronological age is essential when monitoring brain aging not only in health but also in conditions like AD, PD and TBI. Without reference values against which BV changes can be quantified, reliable quantification of disease-related brain tissue loss across adulthood—and especially during old age—is difficult to achieve. For this reason, the availability of a normative reference source which reports human BVs during adulthood is important to both scientists and clinicians. Surprisingly, although many reports of brain volumetrics covering the entire lifespan have been published, high uncertainty remains pertaining to the important relationship between brain volume and age. Firstly, although the temporal trajectory of BV is modulated by sex, many reports have not studied this effect separately for men and women. Secondly, volumetric measurements have traditionally been obtained from a variety of sources, including via MRI tissue classification (Gunning-Dixon et al. 2009; Balafar et al. 2010), computed tomography (CT) image segmentation (Irimia et al. 2019), *post mortem* stereology (Doherty et al. 2000) and using other methods. All these techniques have advantages and drawbacks related to data acquisition and analysis, and all can produce systematic errors of volumetric estimation. Thirdly, brain volumetrics have been reported in a variety of ways, such that the values reported by various studies may be difficult or even impossible to compare. For reasons like these, meta-analyses like the present one are necessary to identify, harmonize and aggregate volumetrics data across the scientific literature. Although many past studies—including some of those included here—provide age-dependent regression coefficients and volumetric estimates for the brain, the key contribution of the present study is its provision of meticulously curated, quantitatively aggregated and rigorously meta-analyzed values for these parameters, with CIs and empirically estimated standard deviations which are considerably more precise than those of any individual study.

### Study selection

In meta-analyses like ours, researchers must pay attention to the eligibility criteria of each study in order to minimize heterogeneity across the aggregate sample of the meta-analysis. For example, many studies have measured BVs and atrophy rates from individuals selected for specific characteristics, whether genetic (e.g. based on their ApoE allele profiles) or environmental (e.g. physical activity levels). Whilst such selection criteria are necessary to study the association between genetics, environment and aging, the meta-analysis researcher must carefully weigh whether—and, if so, how—such studies should be included in a meta-analytic survey of brain atrophy data. If a meta-analysis targeting the general population inadequately aggregates studies whose inclusion criteria result in aggregate samples which do not reflect the typical trajectory of the human adult brain, such an analysis may lead to results and conclusions which do not reflect the characteristics of the normative population whose features it aims to capture. As an illustration, the brain aging trajectory of physically active individuals is likely not very representative of the general population’s overall BV trajectory because activity levels vary considerably from person to person. Furthermore, it may be difficult or impossible to adequately weigh results from different samples based on the extent to which specific sample demographics reflect the composition of the general population. For this reason, studies were excluded from this meta-analysis if their eligibility criteria might have substantially modified the composition of the aggregate sample beyond what is expected from random sampling of the general population.

### Sampling confounds

Because the interactions between genetic, phenotypic and environmental factors upon age-related volumetric trajectories are complex, studies frequently focus on just one of these many contributors which together explain the observed variance of brain trajectories. Some of these variables must be mentioned here due to their importance when interpreting the results of this study. For example, the variation of brain volumetrics with age and sex has been reported extensively by many studies, including those included in this report. Some studies describe brain volumetrics and related measures across the entire range of ages analyzed rather by decade or as a function of age. This can pose challenges to meta-analyses like ours because participants’ age distribution is not always uniform, and the number of subjects whose ages fall within each decade of adulthood is rarely reported. This can preclude the ability to survey the literature in a way which is amenable to reporting volumetric trajectories by decade; one notable exception here is the study of Walhovd et al. (2011), where decadal data are provided. Aside from age, sex is also very important as a biological variable affecting volumetric trajectories (Jancke et al. 2015); for this reason, this study reports regression coefficients and volumetrics separately by sex, where possible.

### Genes and environment

Aside from age, sex and head size, human BV trajectories are partly influenced by genetic and environmental factors whose effects upon brain volume may covary substantially. Lifestyle factors like diet, exercise, intelligence, educational attainment, alcohol consumption and smoking can all affect BV trajectories in a dose-dependent fashion; such factors can be additional sources of variance which is not explained by other factors (Lenroot and Giedd 2008). Genes which modulate the rate of biological aging and human susceptibility to disease can themselves interact with environmental factors. Together, genes, environment and their interaction contribute to brain aging trajectories in complex ways and can account for some variability observed in the measured rates of human brain atrophy (Seshadri et al. 2007). This meta-analysis does not account for the effect of such genetic and environmental variables upon brain trajectories because its purpose is to describe the BV trajectory of the general population rather than of any its subgroups. Nevertheless, future research should aim to meta-analyze the effects of genes and environment upon BV and related measures.

### Allometry

Head size may confound meta-analyses considerably because a structure’s absolute difference in volume over some arbitrary time interval is proportional to the initial volume of that structure. Thus, head size must be accounted for in studies like ours to avoid fallacious inferences driven by effects related to brain allometry (the scaling of the brain with body size) rather than by the main effects of interest, such as aging or disease. Many reports of brain volumes identified in this study did not account for head size or, alternatively, accounted for it in ways which differed enough across studies that case-study comparison was too difficult or, indeed, impossible. Here, regression coefficients were reported as percentages of differences in volume per year precisely to convey information which can be interpreted independently of head size. It should be mentioned that, although normalization by ICV is by far the most common approach to accounting for head size, there is more than one such strategy and certain approaches may be more suitable than others, depending on context (Voevodskaya et al. 2014).

### Volumetric measurement techniques

The descriptive statistics (mean and variance) of brain volumetrics has been made available by a variety of studies utilizing vastly different methodologies. These range from approaches superseded long ago—like post-mortem weighing of fluid volume displaced by the excised brain (Uspenskii 1964)—to state-of-the-art in vivo magnetic resonance imaging (MRI) at ultra-high spatial resolution (Van Leemput et al. 2009). More recently, MRI quantitation methods have been extended to CT (Irimia et al. 2019), thus making BV calculations possible in remote locations or in other environments where MRI acquisition is not feasible (Kaplan et al. 2017). There is typically good agreement between such approaches, even within less than 1% (Despotovic et al. 2015).

This study is restricted to studies undertaken using MRI because this technique currently provides the gold standard for in vivo BV measurement (de Boer et al. 2010). Nevertheless, the potential shortcomings of MRI related to the effects of microstructural properties of brain tissue (e.g. myelination, iron, and water content) upon brain morphology calculations (Lorio et al. 2016; Natu et al. 2019) should be acknowledged. Because the MRI literature is the largest source of brain volumetrics currently available and MRI sample sizes eclipse those of most post-mortem studies very frequently, the MRI literature was found to be more suitable for the present study. An important observation on measurement techniques is that computed volumes may be sensitive to the segmentation method used. Although early studies included here could not benefit from the availability of today’s state-of-the-science MRI analysis software, most reports published after 1999 predominantly used thoroughly validated packages like SPM, FreeSurfer and FSL (Table 4). Usefully, the fact that this meta-analysis relies on volumetrics computed using a variety of segmentation approaches may translate into a reduced probability for its results to be biased toward any single technique. Furthermore, because image analysis methods were treated as covariates in the meta-analysis, their residual confounding effects may not be considerable. Further study, however, is needed to establish the relative prominence of acquisition parameters upon meta-analysis results.

### MRI acquisition parameters

Encouragingly, visual inspection of Figure 1 does not suggest that the regression coefficients and volumetrics reported by older studies differ systematically from those in more recent studies. This, however, does not demonstrate the absence of a relationship between the measures of interest here and the publication dates of the studies included. For this reason, future studies should aim to investigate in more detail the combined effects of MRI spatial resolution, partial volume effects, scanner field strength and of other factors upon the metrics discussed here. The available spatial resolution of anatomic MRI scans has increased considerably over the past thirty years; thus, MRI spatial resolution can confound meta-analyses like ours if their statistical effects are not accounted for. To provide insight on these and other related effects, Table 4 lists key hardware parameters, sequence parameters and analysis methods used in the studies selected for the meta-analysis. For example, whereas early studies relied on both *T*_1_- and *T*_2_-weighted MRI, most recent studies utilized *T*_1_-weighted imaging. More recent studies are typically more likely to provide the opportunities of calculating volumetrics more accurately; currently, the typical voxel size of brain MRI scans is ∼1 mm^3^, which frequently corresponds to a pixel size of 1 mm^2^ and to a slice thickness of 1 mm. Indeed, many studies published in the past ∼15 years and meta-analyzed here feature today’s standard voxel size of 1 mm^3^ or voxel sizes close to it, although this is certainly not the rule (Table 4). It is very important to note that, even when volumetrics are computed from images acquired using the same weighting (e.g. *T*_1_, *T*_2_, or PD), sequence parameters (e.g. *T_E_*, *T_R_*, *T_I_*), spatial resolution and hardware specifications (e.g. scanner field strength) can affect volumetrics substantially even when the same brain segmentation software is used to process data. Partly for reasons like these, the reference data provided in this study should not be interpreted and utilized as highly precise approximations, but rather as statistical estimates provided for the purpose of guidance. As Table 4 illustrates, the variability in sequence parameters across the studies meta-analyzed here is relatively high. Han et al. (2006) have shown that even small variations in sequence parameters (e.g. *T_E_*, *T_R_*, *T_I_*) may affect volumetric calculation results substantially even when images are segmented using the same tissue classification software. Thus, to a large extent, differences between various studies’ computed volumetrics can be due to such methodological differences. Furthermore, the number of excitations (NEX) and flip angle (FA) can influence cortical thickness measurements to a large extent (Han et al. 2006), but the statistical effects of these parameters could not be accounted for in this meta-analysis because they was rarely reported in the original publications. Although a systematic analysis of how sequence parameters affect volumetrics is beyond the scope of this meta-analysis, the literature review leading to the present study appears to convey that one group of critical image acquisition factors affecting the variability of volume calculations include field strength, image weighting and spatial resolution, followed closely by *T_E_*, *T_R_*, *T_I_*, NEX and FA.

In a few older studies included here, the typical spatial resolution is—as expected—poorer than today’s standard. For example, Gur et al. (1991) calculated brain volumes from *T*_2_-weighted MRI in a sample of 69 healthy adults with ages ranging from 18 to 80. Similarly, Blatter et al. (1995) calculated brain volumetrics from *T*_2_-weighted SE MRI from 194 healthy subjects with ages ranging from 16 to 65 years, but their measurements were based on MRI volumes with a slice thickness somewhat greater than today’s standard (Table 4). A third such study is that of Matsumae et al. (1996), who obtained total intracranial volumes from 49 normal volunteers ranging in age from 24 to 80. Importantly, none of these studies were found to yield either cross-sectional regression coefficients or volumetrics which differed in any substantial way from those reported in recent studies (see Table 1 and Figure 1). Furthermore, whereas most older studies had relatively small sample sizes, others were relatively large even by today’s standards (Bigler and Tate 2001) and may therefore still warrant consideration. Thus, Bigler et al. assessed ICV and BV in 532 subjects who were either healthy adults, TBI victims or AD patients. Interestingly, the regression coefficients and volumetrics reported by these older studies did not differ substantially from those reported by recent ones, and the relative sample sizes of the former studies were relatively smaller than those of the latter, such that older studies did not contribute as substantially to meta-analysis results as newer ones did. Additionally, image acquisition parameters were included as covariates in this meta-analysis to reduce their relative effects upon its results. The task of determining how such parameters may individually contribute to volumetric estimates remains challenging and complex because many such parameters affect image quality and volumetrics nonlinearly and are therefore difficult to study using conventional statistical approaches. Nevertheless, it is likely that their inclusion as covariates here partially mitigates their confounding effect upon the measures of interest. As a side note, inspection of Table 1 reveals that the meta-analytic value of *β*^2^ for the entire (aggregate) sample is relatively closer to that of larger, more recent studies than to that of older, smaller studies. This is likely due not only to (A) greater effects due to larger studies, and also to (B) the correction for the confounding effects of acquisition parameters (e.g. voxel size). Since spatial resolution is higher in more recent studies, it is conceivable that accounting for spatial resolution confounds and related effects may have additionally altered the contribution of older studies to the meta-analysis compared to that of more recent studies.

### Utility of reported measures

Frequently, researchers who utilize brain volumetric data are interested in two important groups of measures: (1) the BV trajectory as a function of age, and (2) the mean and variance of BV at a given age and/or for each sex. The first such group of parameters includes linear regression coefficients describing the relationship between volumetrics and age across the range from 20 to 70 years. The second group includes average volumetrics at selected ages (20 years, 45 years, and 70 years). Researchers have many choices as to the nature, format and units used to report such data on the relationship between brain volumetrics and age. Depending on the context in which such values are required, certain choices may be preferable to others depending on the researcher’s objective needs and/or subjective preferences. In this study, regression coefficients are reported as percentages of differences in volume per year, rather than in cm^3^ per year, ml per year or in other units. Reporting coefficients using *percentages* rather than *physical units* of volume has a considerable advantage, in that the former choice is independent of head size whereas the latter is not. Thus, a regression coefficient reported as a percentage difference in volume per year does not suffer from the confounding effect of varying head sizes, either within or between studies. Nevertheless, because scientists frequently need to calculate the expected values of physical volumes based on regression coefficient data, it is also important to have access to reference values of average volumetrics for the end points of the age interval over which regression coefficients were computed (20 to 70 years in the present study). This is one of the reasons for which this study reports average volumetrics for men and women at the ages of 20, 45 and 70, which correspond conveniently to representative—albeit somewhat arbitrary—values for young, middle, and old age.

Importantly, availability of the parameters reported by this meta-analysis allows one to estimate the average value of volumetrics at *any* age of interest within the stated interval from 20 to 70, under the assumption that age and volume are linearly or quadratically related. Thus, knowledge of these two important sets of parameters can facilitate the process of obtaining reference values for brain volumetrics at any adult age of interest within the range considered. Specifically, based on the linear model, one can calculate the expected value of volume *V* at age *t* within the range from 20 to 70 by (A) identifying the value of the desired volume *V* at age 20 in Table 3, and then (B) calculating the expected value of *V* at time *t* (age in years) as

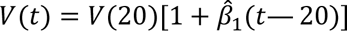

where 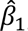 is the corresponding regression coefficient estimate for the volumetric measure of interest, as reported in Table 1. For example, to calculate females’ average BV at age 60, one can proceed by identifying the average of *V*(20) in Table 3 (i.e. 1,274 cm^3^) and then use 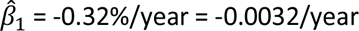 from Table 1 to calculate an approximate value of 1,111 cm^3^ for the average female BV at age 60. Thus, one strength of calculating average brain volumes from meta-analysis results like ours is that meta-analyses aggregate data over a range of studies, presumably yielding more accurate estimates.

An important benefit of the present meta-analysis is that it can assist researchers to evaluate the relative standing of their own volumetric analysis results relative to other results reported in the literature. Specifically, because this study reports first-order statistics and confidence intervals, such information can be used by scientists to calculate the extent to which the means and standard deviations of their own volumetrics deviate from the aggregate sample in the meta-analysis. This can be accomplished, for example, by using Hotelling’s *T*^2^ test to estimate how different a dataset’s mean volumetrics are from the means of the aggregate sample studied here. Alternatively, researchers can use the tables for reference to determine which sample(s) is/are most suitable for comparison to their own datasets; then, a researcher can utilize Hotelling’s *T*^2^ test or other suitable statistical tests to compare her/his sample to the reduced reference sample deemed to be most adequate for such comparison.

Aside from expediting the calculation of average volumes, the results presented here also facilitate comparisons between health and disease. For example, Cole et al. (2018) found that, annually, victims of moderate-to-severe TBI may experience, on average, a mean GMV loss of 1.55%, a mean WMV loss of 1.49% and a mean BV loss of 1.51%. Comparison of these annual volume differences with those listed for in Table 1 healthy adults clearly supports the conclusion of Cole et al. that TBI patients experience GMV, WMV and BV trajectories which are substantially steeper than those observed in typical aging. Similar comparisons can be made for other neurological conditions, like AD and PD.

### Cross-sectional vs. longitudinal studies

Cross-sectional studies suffer from the major limitation that they only allow the investigation of age-related *differences* and *trajectories* rather than changes, partly because of cohort effects, secular trends and uncontrolled individual differences (Raz and Rodrigue 2006). The results of the present study should therefore be interpreted with awareness of this important caveat. By contrast, longitudinal studies of brain volumetrics can be more valuable because they allow researchers to estimate brain atrophy rates within specific individuals and across specific time periods while (partially) controlling for individual differences.

Unfortunately, the literature survey conducted identified relatively few longitudinal studies of brain atrophy in youth or middle age compared to old age. For this reason, data on BV trajectories across adulthood are more abundantly available from cross-sectional—rather than longitudinal—studies. Additionally, many—if not most—longitudinal studies of brain atrophy focus on comparisons between groups whose membership is limited by relatively narrow eligibility criteria. Such criteria can encumber the inclusion of longitudinal studies in meta-analyses because different eligibility criteria may not only confound the meta-analysis, but also result in an aggregate sample whose characteristics may not be adequately representative of the general population. As importantly, few longitudinal MRI studies have measured brain volumetrics over periods exceeding 10 years and none have followed up participants over the entire course of their adulthood. This is partly due to the substantial practical challenges of doing so, and to the fact that MRI and related tools for the calculation of brain volumetrics have been widely available for only ∼30 years. Incidentally, two other potential drawbacks of longitudinal studies are survivorship bias (Nunney 1991) and self-selection bias (Heckman 1990). Thus, despite the many advantages of longitudinal studies, there is still considerable appeal in relying on cross-sectional brain volumetrics data when characterizing BV trajectories throughout adulthood. The reader is referred to the contributions of Raz & Rodrigue (2006) for further discussion of the relative merits of cross-sectional vs. longitudinal studies of brain morphometry.

### Failure to report total volumes

Whereas most early MRI studies quantified only total volumes, many studies from the past decade have shifted focus to the detailed reporting of regional volumes and atrophy rates. One unfortunate and paradoxical side effect of this trend toward greater regional specificity and sophistication is that the total volumes of GM, WM and CSF are no longer being reported as often as before the advent of regional volumetric analysis. This can have unfortunate consequences because brain parcellation schemes often differ considerably across studies, such that various brain regions’ measures fail to be reported in a way which facilitates summary. Thus, it may be tedious or impossible to calculate total volumes from regional volume data unless these data are complete and the parcellation schemes employed are specified unambiguously. Furthermore, many studies leveraging data from large repositories and efforts such as the Alzheimer’s Disease Neuroimaging Initiative (ADNI, http://adni.loni.usc.edu), the Parkinson Progression Marker Initiative (PPMI, https://www.ppmi.info.org) and the UK Biobank (https://www.ukbiobank.ac.uk) may fail to report the total brain volumetrics of healthy control subjects included in these studies; alternatively, their reported volumetrics may apply to healthy control individuals whose inclusion criteria and demographics may differ substantially across any pair of datasets from these neuroimaging consortia. For reasons like these, it is very important that future studies continue to report BV, GMV, WMV and CSFV in addition to regional volumes.

### Linear vs. quadratic models

Many studies of brain volumetrics—whether longitudinal or cross-sectional— have modelled the time-dependent trajectories of brain volumetrics using either linear or quadratic regression models. In a strict quantitative sense, quadratic fits are better than linear ones because the former yield a smaller sum of squared residuals, i.e. better goodness of fit. Nevertheless, due to the danger of overfitting in studies like ours, the question as to which model is more justified empirically deserves discussion. In some studies included here—e.g. Taki et al. (2004)— linear fits seem appropriate, and the quadratic relationship between volumetrics and age does not appear to be substantial. In others—e.g. Sowell et al. (2003)—visual inspection suggests that the contribution of the quadratic term to the BV trajectory is important. It has been proposed that the choice between linear and quadratic models is largely dependent upon the age range of participants in a given sample, and that researchers should strive to explore as broad a range as possible (Fjell et al. 2010). Furthermore, the inclusion of participants in the first and second decades of life may strengthen the argument in favor of using quadratic models due to the trajectory of brain dynamics during development (Sowell et al. 2003). The present meta-analysis of volumetric trajectories across adulthood provides evidence in support of the assertion by Fjell et al. (2010) according to which the most seemingly appropriate fit is determined by the range of ages included. Good et al. (2001) did not find significant WM decline with age in their sample of 465 healthy adults; these authors suggested that, in the first six decades of life, aging affects GMV more than WMV, and that both GMV and WMV decrease substantially after age ∼60. Thus, the quadratic regression curves depicted in these authors’ study suggest some GMV increase up to the fifth decade of life, with relatively slow subsequent decline into the eighth decade; by contrast, the decrease in WMV appears to be monotonic. These trajectories are reproduced by other studies (Fotenos et al. 2005; Walhovd et al. 2011). In particular, the results of Walhovd et al. (2011) indicate that cerebral GMV and BV decrease monotonically throughout adulthood, suggesting that a linear fit may be appropriate for these measures. By contrast, cerebral WMV exhibits nonlinear behavior in that it increases slightly between ages ∼20 and ∼60, and then decreases relatively faster than within this interval. Figure 2 can assist in assessing the relative merits of linear and quadratic models by reproducing their predicted trajectories. These depictions, however, should be interpreted carefully because the choice between a linear and a quadratic model may depend substantially both on the population from which samples are obtained and on the sizes of these samples.

The model selection results based on the AIC analysis suggest that, in the case of BV and CSFV, the quadratic model is only marginally better than the linear model at minimizing information loss. For GMV, the quadratic model is of somewhat greater merit, whereas for WMV the benefit of the quadratic model is substantial. The present meta-analysis found that, historically, studies based on quadratic models have been far less common than those using linear models. Because the eligible quadratic model studies identified here were few and had a relatively modest combined sample size, future studies should explore the suitability of quadratic models in further detail to warrant more rigorous meta-analysis. This argument is strengthened by the results of the AIC-based analysis pertaining to the relative information loss between linear and quadratic models.

### Meta-analytic limitations

It is important to acknowledge that this meta-analysis is neither exhaustive nor definitive. For example, although many studies were reviewed to generate it, it is both plausible and likely that many studies which fit the selection criteria of the meta-analysis were not included because of missed opportunities to identify them. Furthermore, different study selection criteria may have resulted in rather different meta-analysis results, and it is not clear that the selection of studies included here is the best possible selection. For example, studies published before 1990 were not included and very recent studies may not have been available in the public literature databases consulted at the time when the study was undertaken. For some measures, the lack of overlap between the CIs of several studies (see Figure 1) may suggest that some sources of between-study heterogeneity could not be removed even by implementing the stringent search criteria of the meta-analysis. It is also possible that some metrics included in the meta-analysis were systematically affected by confounds which were not discussed here. Finally, the values of various parameters extracted from each study may not be as numerically accurate as those originally computed by the respective study’s authors based on their original data. All these limitations of this analysis are duly acknowledged. Nevertheless, this meta-analysis does cover a relatively large portion of the scientific literature on brain volumetrics and therefore has the potential to be of substantial utility to scientists and clinicians.

## Conclusion

BV trajectories in adulthood are of substantial interest in studies of human aging. Historically, regression coefficients describing the relationship between age, BV and related metrics have been under-reported by MRI studies of human volumetry despite many researchers’ need to compare brain trajectories across different populations to map effects associated with disease, genetics or environmental factors. Similarly, the average volume of the brain is a key fact about humans which is illustrative of mankind’s uniqueness, and the ability to calculate its value at various ages during adulthood based on a normative, aggregate sample is very important to neuroscientists, gerontologists, anthropologists and clinicians. The cross-sectional data provided in this meta-analysis can facilitate such calculations and allow researchers to gain further insight to assist the interpretation and assessment of the literature on this important topic. Nevertheless, readers should exercise caution when using and interpreting the meta-analytic estimates provided in this study due to their dependence on MRI hardware specifications (e.g. scanner manufacturer, magnetic field strength), data acquisition parameters (e.g. spatial resolution, weighting), and brain segmentation methods. Future studies of brain volumetrics across the lifespan should strive to provide decadal data and to explore the nonlinearities of age-dependent BV trajectories. Because there are relatively few longitudinal reports of brain atrophy during youth and middle age, more such studies should be undertaken to quantify BV loss. This could facilitate and advance the necessary transition from surmising volumetric trajectories from cross-sectional data to their fiducial estimation from longitudinal studies.

## Conflict of interest statement

The author declares that he has no conflict of interest.

## Acknowledgments

This work was supported by the National Institutes of Health under Award Numbers RF1 AG 054442 and R01 NS 100973, and by the US Department of Defense under contract W81XWH-18-1-0413. The content is solely the responsibility of the author and does not necessarily represent the official views of the National Institutes of Health or of the Department of Defense.

## Abbreviations

AD: Alzheimer’s disease
AIC: Akaike information criterion
AIR: automated image registration B magnetic field strength
BET: brain extraction tool
BV: brain volume
CSF: cerebrospinal fluid
CSFV: erebrospinal fluid volume
CT: computed tomography
EM: expectation maximization
ETL: echo train length
ExBrain: extended brain
F: female(s)
FSL: FMRIB software library
GE: General Electric
GM: gray matter
GMV: gray matter volume
ICV: intracranial volume
M: male(s)
MIDAS: metabolite imaging and data analysis system
mm: millimeter
ms: millisecond
MRI: magnetic resonance imaging
PD: Parkinson’s disease
RAVENS: regional analysis of volumes examined in normalized space
SPM: statistical parametric mapping
T: tesla
TE: echo time
TI: inversion time
TR: repetition time
TBI: traumatic brain injury var various
VBM: voxel-based morphometry
WM: white matter
WMV: white matter volume

## Notes

### Competing Interest Statement

The authors have declared no competing interest.

### Summary of Updates

This revision is an update which reflects the accepted manuscript. It contains modifications, corrections and additions to the main text, figures and tables.

